# Early life sleep is associated with longevity in *Drosophila*

**DOI:** 10.64898/2025.12.05.692620

**Authors:** Eshani Yeragi, Elizabeth B. Brown, Jessica A. Bernard, Alex C. Keene

## Abstract

Sleep is a universal behavior that is critical for brain function and physiological homeostasis. While growing epidemiological and experimental evidence suggests reduced sleep quality is associated with negative health outcomes, the causal relationship between sleep loss and reduced longevity remains poorly understood. Here, we examine sleep across the lifespan and its relationship to longevity in *Drosophila melanogaster*. We examined the associations between numerous components of sleep at different life stages to longevity. Early life sleep, but not middle and late-life sleep, was positively associated with longevity. Age-dependent changes in sleep were consistent but accelerated, when flies are housed under stressful conditions including nutrient and temperature stress. Further, pharmacological restoration of sleep in the first 10 days of life, but not at later time points, increases longevity. Together, this work provides a systematic investigation of how different components of sleep impact longevity and suggests early life sleep may be particularly important in promoting sleep across the lifespan.

## Introduction

Disorders that impair sleep duration and quality are associated with severe negative health outcomes, including reduced cognitive function and longevity [1] . While humans are often thought to need ∼8 hours of sleep per day, the duration of sleep varies significantly between individuals, as well as within individuals across their lifespan [2]. Sleep duration in humans is highest in early life dropping during young adulthood and lowest in midlife, while sleep quality diminishes in later life[3], and is modulated by environmental variables including stress or social context [4–7]. Because sleep is influenced by many environmental and developmental factors, understanding how variation in genetic architecture contributes to sleep regulations across contexts is a challenge for sleep medicine.

Large scale human Genome Wide Association Studies (GWAS) and public health surveys have sought to examine the relationship between genetic variation and contextual regulation of sleep[8,9]. Most of these studies rely on self-reporting, while long-term actigraphy studies are typically limited in size[10,11]. Together, these analyses have identified numerous associations between sleep and environment including dietary habits, ambient temperature, physical activity, stimulant use, and screen exposure before bedtime, all of which significantly influence sleep duration, quality, and circadian regulation [12]. Furthermore, the co-morbidity between sleep loss and many other diseases, as well as the age-dependent loss of sleep duration and quality, reveal a critical role for sleep in healthy aging[13,14]. Developing an animal model to test the relationship between sleep and aging allows for systematic analysis of the relationship between sleep and aging.

The fruit fly, *Drosophila melanogaster,* is a leading model for studying genetic regulation of sleep and aging [15,16]. The mechanisms underlying each of these processes are highly conserved from flies to mammals[17]. Flies, like mammals exhibit distinct electrophysiological patterns that correlate with wake and rest [18,19], and we have identified sleep-associated reductions in metabolic rate in flies that are consistent with those that occur in mammals [20–22]. In addition, flies display all the behavioral hallmarks of sleep, including an extended period of behavioral quiescence, rebound following deprivation, increased arousal threshold, and species-specific posture [23,24]. Chronic sleep deprivation or circadian disruption results in shortened lifespan in *Drosophila,* suggesting a critical role in modulating health. Further, in flies, like in humans, sleep deteriorates in later life, revealing conserved age-related changes in sleep are shared between invertebrates and mammals [25].

The quality and duration of sleep in *Drosophila* changes over the lifespan of the fly [26,27]. Sleep duration is greater early in life, and this is critical for neurodevelopment [28,29]. As flies age, sleep becomes more fragmented, indicating reduced sleep quality[26]. Further, expression of Alzheimer’s associated genes, or the pharmacological induction of reactive oxygen species accelerates age-related sleep fragmentation [26,30]. Age related changes in sleep are also impacted by diet, social context, and early life experience [31,32]. Dietary, genetic and pharmacological screens have identified factors associated with age-related changes in sleep [33], yet the impact of naturally occurring variation in sleep across the lifespan and longevity remains largely unexplored.

Here, we sought to measure sleep across the lifespan, and its relation to longevity in individual *Drosophila melanogaster*. Lifelong analysis of sleep in individual flies provided the opportunity to examine how different variables of sleep, including duration, bout length, bout number, and circadian regulation of sleep, change across the lifespan, and how this impacts longevity. These studies identified a critical association between early life sleep and longevity across multiple populations.

## Results

To examine how sleep changes across the lifespan, we measured sleep in *w^1118^* flies, a commonly used genetic background for genetic experimentation[34–36]. Female flies were placed in Drosophila Activity Monitors (DAMS) to record sleep and activity three days following eclosion and maintained in tubes throughout their lifetime [23,24]. Like previous studies, flies were transferred to fresh food every five days for the duration of their lifespan [26,37]. Quantifying the longevity of flies under these conditions revealed *w^1118^* flies had a median survival of approximately 59 days (Fig. 1A). This lifespan is consistent with findings from other groups that have measured survival in group-housed flies or in vials offering a larger habitat, suggesting that the protocols used here do not substantially impact lifespan [38,39]. Based on this longevity analysis, we defined young flies as (4-14 days), middle life (days 24–34) and in old age (days 54–65).

**Figure 1.**
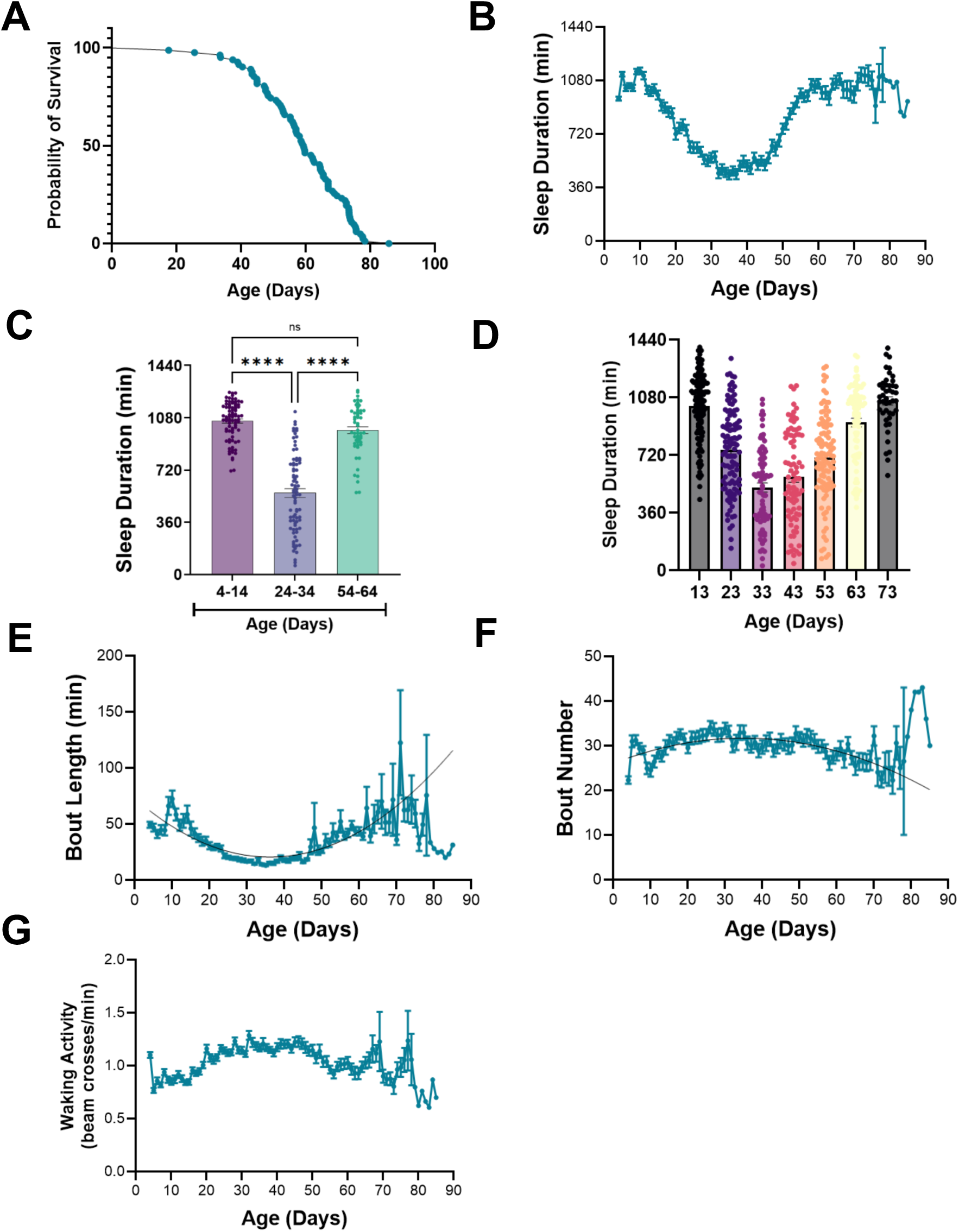
Changes in sleep across lifespan. 1A) Kaplan–Meier survival analysis of individually housed *w^1118^* flies on the standard food showed mean survival of 58.85 days. 1B) Average daily sleep duration (in minutes) measured across lifespan of *w^1118^* flies exhibited a U-shaped trajectory with an early-life decline in sleep followed by recovery of sleep by mid-to-late life *(R^2^=0.3009)*. 1C) Sleep duration varied significantly across three distinct life-stages (mixed-effects repeated measures ANOVA*: F (1.384, 92.05) = 160.1, p < 0.001*). Tukey’s post-hoc test exhibited a decline in sleep duration from Day 4-14 to Day 24-34 followed by recovery in late life Day 54-64 *(p < 0.001)* with Two-way ANOVA confirming age as a significant factor (*F (2, 133) = 162.2, p < 0.001*). 1D) A two-way ANOVA revealed a significant effect of age (*F (6, 491) = 57.39,* p < 0.001) in sleep duration across seven distinct ages in socially housed *w^1118^* flies similar to the pattern observed in individually housed flies. Tukey’s comparisons showed reduced sleep-in mid-life (13day vs. 33day: *p<0.001*), with recovery in late life (53day vs. 73day: *p<0.001*) with no significant difference in sleep duration across 13day vs. 73day (p>0.99). 1E-F) Bout length was greater in early and late life, reflecting consolidated sleep (*R^2^=0.08*) with U-shaped trajectory across the lifespan, whereas bout number showed an inverted U-shaped pattern across the lifespan with fewer bout numbers at lifespan. 1G)

We quantified sleep across the lifespan using the standard metric of 5 minutes of immobility to define sleep[24]. Sleep duration was highest in early life (days 4–14), declined during mid-life (days 24–34), and increased again in old age (days 54–65) (Fig. 1B, C). These age-dependent changes were evident during both the daytime and nighttime, indicating that alterations in sleep duration occur across the entire circadian cycle rather than being restricted to a specific time of day (Fig. S1A, B). Together, these results demonstrate robust, stage-specific changes in sleep across the lifespan.

Sleep, like many behaviors, is modulated by social context [6,40]. Because our analysis protocol involves individually housed flies, it raises the possibility that long-term social isolation could contribute to age-dependent changes in sleep. To directly test this, we aged *w*^1118^ flies under group housed conditions and transferred them to activity monitors at defined time points. These flies resembled individually housed flies, with high sleep levels in early life, reduced sleep-in mid-life, and a subsequent increase in late life (Fig. 1D). Thus, the observed lifespan changes in sleep are not attributable to long-term social isolation.

Total sleep duration is regulated by both the frequency and duration of individual sleep bouts and can be modified through changes in either or both parameters. Regression analysis reveals that both bout length and bout number vary significantly across the lifespan (Fig. 1E, F), suggesting changes in both these variables contribute to changes in sleep duration.

The application of Hidden Markov Models has been used to estimate the probability of maintaining a waking state p(Wake) or a sleeping state p(Doze), the latter proposed to reflect sleep depth. We performed analysis on the same age periods across the lifespan and found that p(Wake) was elevated in midlife, while p(Doze) was reduced (Fig. S1C,D,E), supporting the notion that sleep need is reduced in *w^1118^* flies at midlife. Two-way ANOVA revealed a significant main effect of age but no independent effect of state (pWake vs pDoze). Post-hoc tests showed that early-versus mid-life and mid-versus late-life periods differ significantly, whereas early and late life did not. To determine if activity differs across the lifespan, we measured waking activity, normalizing the amount of total activity for time awake. Waking activity increased in midlife, inversely correlating with changes in bout length and sleep duration (Fig. 1G). Therefore, in *w^1118^* flies display robust changes in sleep duration, sleep architecture, and activity over the lifespan.

To identify potential interactions between sleep and longevity, we quantified the relationship between individual sleep, activity, and longevity variables. Specifically, we compared sleep duration, bout number, bout length, and waking activity during day and night periods in relation to longevity during the day and night. Each trait was measured during early life (Day 4–14), mid-life (Day 24–34), and late life (Day 54–64). For early-life measurements, total sleep duration, day and nighttime sleep duration were positively correlated with longevity (Fig 2A). Furthermore, bout measures of increased sleep depth including bout length and p(Doze) were also positively associated with increased longevity (Fig 2A). Conversely, bout number and p(Wake), two factors linked to reduced sleep quality were negatively associated with longevity (Fig 2A). Together, these findings support the notion that sleep duration, and the consolidation of sleep during early life promotes longevity.

**Figure 2.**
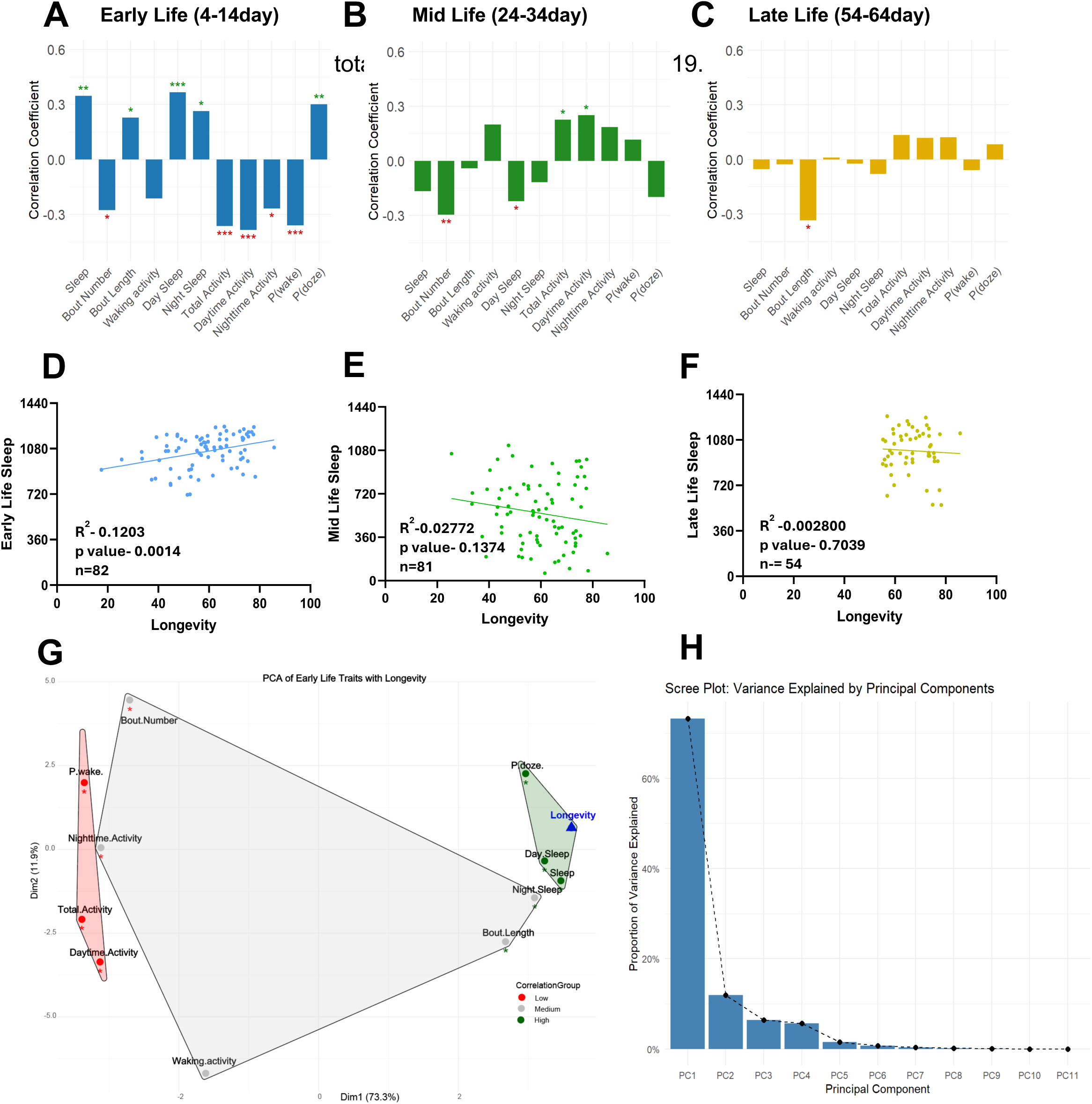
Associations between sleep traits and lifespan. 2A) Early Life (4-14day) bar correlation analysis exhibited total sleep (*r = 0.35, p < 0.01*), day sleep (*r = 0.37, p<0.001*), night sleep (*r = 0.26, p<0.05*), and bout length (*r = 0.23, p<0.05*) to have positive correlations with lifespan. Daytime activity (*r = −0.39, p < 0.001*), nighttime activity (*r = −0.27, p<0.05*), total activity (*r = −0.36, p<0.001*), and bout number (*r = −0.28, p<0.05*) showed negative correlations. 2B) Mid-life trait correlations to longevity revealed bout number (*r = −0.30, p<0.01*) and day sleep (*r = −0.22, p<0.05*) to have significant negative correlations. Total activity (*r = 0.23, p<0.05*) and daytime activity (*r = 0.25, p<0.05*) showed significant positive correlations. Other traits did not reach significance (*p>0.05*). 2C) Late-life traits showed no significant associations with longevity. Only bout length was significantly correlated (*r = −0.33, p < 0.05*). All other sleep and activity traits were non-significant (*p>0.05*). 2D-F) Early-life sleep showed a significant positive relationship to longevity (*R² = 0.12, p<0.01*), whereas regression of mid-life *(R² = 0.03, p>0.05)* and late-life (*R²<0.01, p>0.05*) total sleep versus lifespan had no significant association detected. 2G-H) PCA biplot of early-life traits showed Total sleep, day sleep, night sleep, bout length, and P(doze) to be on the positive side of PC1, while activity-related traits like total, daytime, nighttime activity; waking activity; P(wake) were on the negative side. Scree plot revealed that PC1 and PC2 together explained 63% of variance in early-life trait structure (PC1 = 43.1%, PC2 = 19.9%). 2A–2C show Pearson correlation

Unlike the findings in early life flies, there was no correlation between sleep and longevity in mid and late-life flies (Fig 2B, C). Interestingly, total activity, as well as nighttime and daytime activity, was negatively correlated with longevity during early life but positively correlated during midlife (Fig. 2A-C). Furthermore, bout number was negatively associated with longevity in mid-life, while bout length was negatively associated with longevity during late life (Fig. 2B-C). Together, these results support the notion that the effects of sleep on longevity changes across the lifespan and raise the possibility that the function of sleep also changes across this period.

To further quantify the relationship between sleep and longevity, we performed a linear regression analysis in early, mid and late-life flies. This analysis revealed that early-life total sleep explained 12% of the variance in longevity whereas there was no significant relationship in mid-life and late-life sleep (Fig 2D-F). Additionally, we conducted a principal component analysis (PCA) incorporating all measured early-life sleep parameters (Fig. 2G–H). The scree plot (Fig. 2H) shows that the first two principal components together accounted for 63% of the total variance (PC1 43.1%, PC2 19.9%), suggesting that two components explain the major sources of variation. Traits on the positive side of PC1 follow the same trend, whereas traits on the negative side trend in the opposite direction. Total sleep, bout length, day sleep was positioned on the strongest positive side of PC1, aligning their significant positive correlations with longevity. Conversely, traits such as total activity and daytime activity loaded negatively on PC1, consistent with their negative correlations with lifespan. The convex hull clusters in the PCA biplot further illustrate that traits promoting consolidated sleep (longer bout length, higher total sleep and P(doze)) group together with higher longevity correlations, while fragmentation and high waking activity cluster with reduced lifespan. Together, this multivariate pattern underscores that early-life sleep architecture accounts for a significant portion of lifespan variability, emphasizing the importance of early sleep quality in shaping healthy aging trajectories.

The first two principal components from mid-life trait analysis explained 63% of total variance (PC1 46%, PC2 17%) (Fig. S2A). Sleep-related traits loaded positively on PC1, including total sleep, day and night sleep, P(doze), and bout length, whereas activity variables such as total activity, daytime and nighttime activity, waking activity, and P(wake) loaded in the opposite direction. Mid-life PC scores showed a positive association with lifespan, although the relationship was noticeably weaker than what we observed during early adult life (Fig. S2B). In late life, PCA accounted for less of the overall variance (PC1 42%, PC2 17%) (Fig. S2C). The analysis showed that bout length and bout number contributed the most to PC1, but longevity did not align clearly with either sleep or with activity traits (Fig. S2D). Together, these results show that early-life sleep duration serves as the most effective predictor of lifespan, mid-life sleep traits retain some predictive value but less so, and late-life traits fail to explain sleep-longevity relationships.

Various life history factors including diet and temperature influence sleep [41,42] . It is possible that the relationship between sleep and longevity persists across environmental conditions, or that they are limited to standardized conditions when animals are exposed to limited stress. We sought to determine whether the relationship between early life sleep and stress persists in animals exposed to long-term environmental stress. Nutritional stress was induced by housing flies on a diet composed of only 10% sucrose starting at 4 days old. A separate group of flies was exposed to temperature stress by maintaining them at 28°C from day four onward (Fig 3A). Control flies were maintained at 25°C on standard food throughout their lifespan. Both nutritional stress and temperature stress significantly shortened the lifespan compared to control flies (Fig 3B). In both the temperature and nutritional stress groups, the ontogenic changes in behavior including a reduction in sleep followed by late-life increase that appeared intact yet accelerated, with the notable difference that early life sleep was reduced in the nutritional stress group (Fig 3C).These same patterns of accelerated aging were apparent with changes in both length and bout number (Fig S3A, B). Waking activity also differed

**Figure 3.**
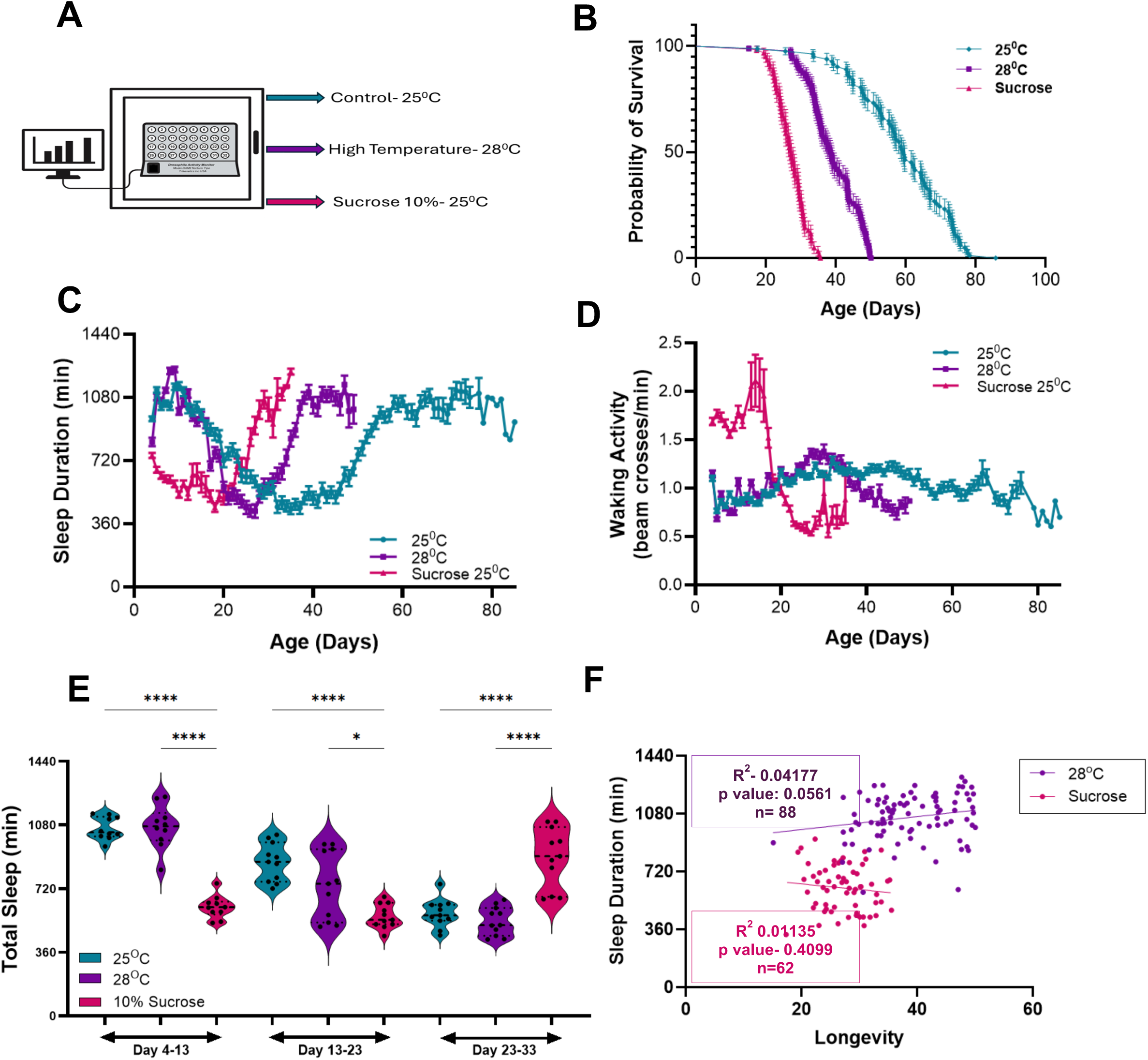
Effects of environmental stressors on sleep profile across lifespan. 3A) Diagram depicts rearing conditions: 25°C (control, black), 28°C (thermal stress, purple), and 25°C with sucrose-only diet (dietary stress, pink), across the lifespan. 3B) The high temperature and the sucrose group showed significantly shorter survival compared to the control group with sucrose being most detrimental over three (Log-rank (Mantel–Cox) test: *χ² = 322.1, df = 2, p<0.001*). 3C) Sleep duration and trend remained similar to the controls, however there seemed decrease in early life sleep in sucrose group The mean sleep remained significantly different across the groups (One-way ANOVA: *F(2,156) = 3.227, p<0.05, R²=0.04*). 3D) One-way ANOVA showed a significant difference in waking activity between groups (*F(2,154) = 4.13, p<0.05; R² = 0.05*). Tukey’s post hoc test showed higher waking activity in the sucrose group compared with 25 °C and 28 °C (p<0.05). There was no significant difference between 25 °C and 28 °C (*p>0.9*). 3E) There was significant effect of stress on total sleep (*F*(8,77) = 33.02, *p*<0.001) with no significant age-stress interaction (*F(10,77) = 0.391, p>0.9).* Dietary stress significantly reduced sleep in early life (*p<0.001*), whereas there was significant effect of thermal stress in mid-to-late life (*p<0.001*). 3F) Linear regression analysis showed no significant correlation between early-life sleep and longevity under thermal and dietary stress (28°C- *R² = 0.04, p>0.05*, 10% sucrose - *R² = 0.01, p>0.4*).

To account for shorter lifespans of flies under stressful conditions, we quantified sleep across three life stages: early life (days 4–13), mid-life (days 13–23), and late life (days 23–33). To determine whether the relationship between early-life sleep and longevity persists under stress, we assessed the correlation between early-life sleep and lifespan in flies housed on different environmental stressors. Flies subjected to nutritional stress slept significantly less during early life and significantly more in late life compared to those under temperature stress or control conditions. In contrast, sleep patterns in flies exposed to temperature stress did not differ from controls at any time point (Fig. 3E). Regression analysis of longevity to sleep duration showed no significant association under either stressors (Fig. 3F). These findings demonstrate that nutritional stress has distinct, long-lasting effects on sleep and activity that are not observed with temperature stress.

We sought to determine whether the relationship between early-life sleep and longevity persists in flies exposed to stressful conditions. Regression analysis revealed a relationship between early life sleep and longevity in the temperature stress group, phenocopying control flies (Fig. 3F). Conversely, Principal component analysis (PCA) further supported this finding; the association between sleep and longevity was nearly absent in flies maintained on a high-sucrose diet, while sleep duration remained the most closely associated with lifespan under temperature stress (Fig S3C-F). Under high temperatures the early life traits had the same relationship to longevity as with controls. Early life traits on high sucrose diet had no distinct association to longevity; in contrast, mid-life traits under both high temperature and high sucrose diet showed distinct reversal, where sleep traits were linked to short life-span, while activity related traits and P(wake) were associated to increase longevity. In early life under both normal and high temperature conditions it showed higher bout length and lower bout number which suggests consolidated sleep was linked to higher longevity. However, this association was absent in early-life high sucrose diet group. In mid-life under both stressors there was reversal in the pattern where longer bout length was associated with low longevity. Together, these findings suggest that physiological stressors, particularly nutritional stress, disrupt the relationship between early-life sleep and longevity.

The association between early life sleep and longevity raises the possibility that increasing early life sleep can extend lifespan. In *Drosophila*, the GABA agonist gaboxadol increases sleep in young and old flies [43,44]. To determine whether increase sleep extends lifespan, we housed flies on food containing gaboxadol from days 3-13, 13-23, or 23-33 (Fig 4A). In agreement with previous findings, gaboxadol administration significantly increased sleep at all measured time points relative to controls. Sleep levels returned to baseline within 10 days after flies were transferred back to standard food. (Fig 4B, C) [43]. Analysis of the effects of gaboxadol on 10 days following the cessation of treatment reveals that sleep is significantly lower than during treatment across all three timepoints (Fig. 4B and S4C). Gaboxadol treatment also impacted on key parameters of sleep including bout length, bout number, p(Wake) and p(Doze) and returned to normal levels following the cessation of treatment (Fig S4A-H). Therefore, gaboxadol provides a method to reversibly enhance sleep across defined timepoints.

**Figure 4.**
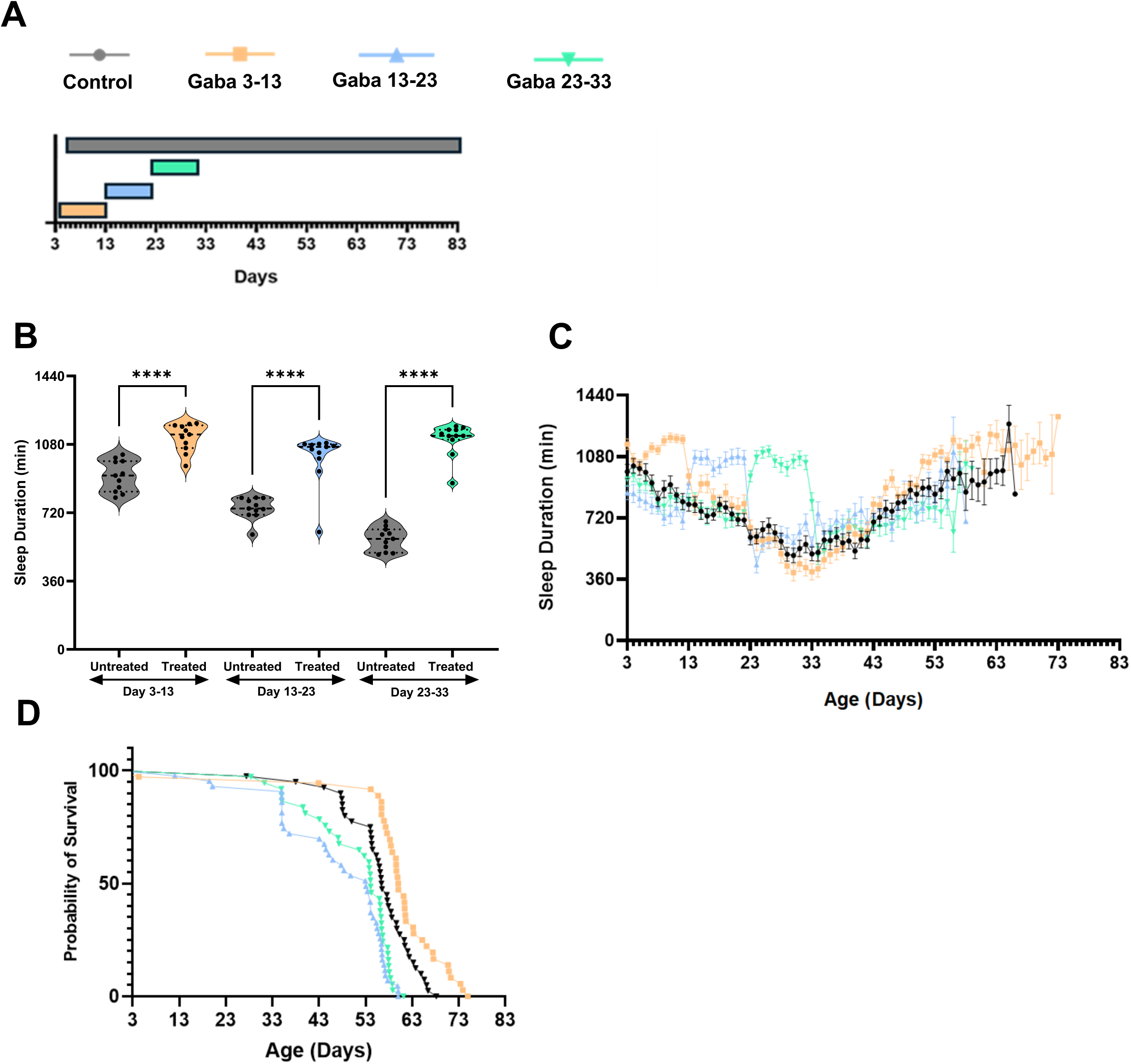
Early life sleep increases longevity. 4A) Four groups were examined: an untreated control group (black) maintained on standard food throughout lifespan, and three treatment groups exposed to Gaboxadol during specific 10-day period—early-life (Day 3–13, orange), mid-life (Day 13–23, blue), and late-life (Day 23–33, green). 4B) Gaboxadol significantly increased sleep across all treatment windows compared to untreated controls (*p < 0.001*). 4C) Two-way ANOVA revealed significant main effects of age (time) and treatment group (*p < 0.001*) as compared to the control. Dunnett’s post hoc comparisons indicated that all Gaboxadol-treated groups exhibited significantly increased sleep relative to controls (*p < 0.001*), with the most pronounced and sustained effect observed following early-life treatment (Day 3–13). 4D) Gaboxadol treated group exhibited increase in longevity as compared to the control and untreated groups. Log-rank (Mantel–Cox) test showed a significant difference among groups (*χ² = 63.39, df = 3, p< 0.001*). Bars represent mean ± SEM with individual points.

Next, we sought to examine the relationship between gaboxadol-enhancement at defined timepoints and longevity. Flies fed gaboxadol from days 3–13 exhibited significantly longer lifespans than those fed gaboxadol from days 13–23, 23–33, reveling a critical window for the effects of sleep on longevity (Fig 4D). In addition, flies treated with gaboxadol between from days 3-13 lived longer than control flies. Together, these findings reveal a critical window where enhancement of early-life sleep promotes longevity.

## Discussion

Studies in humans and model organisms have examined the associations between various life-history traits and lifespan [45–47]. However, this approach is particularly challenging in long-lived species, where longitudinal analyses are often impractical, due to statistical power and challenges controlling key environmental variables[48]. A key advantage of using *Drosophila melanogaster* is its short lifespan, which enables continuous behavioral monitoring across the entire life of the organism [16]. This makes it possible to apply longitudinal methodologies commonly used in the public health field to study how ontogenic and age-related changes in specific traits relate to lifespan outcomes. In our study, we found that several sleep-related measures, including daytime and nighttime sleep duration, as well as statistically modeled sleep depth, were positively associated with increased longevity. Importantly, these relationships were only evident early in life, with no detectable association between sleep metrics and longevity in later life, underscoring the temporal specificity of trait–lifespan associations. These findings illustrate the power of *Drosophila* as a model for dissecting the dynamic relationships between behavior and health across the full lifespan.

Here, we examined sleep across the lifetime in isogenic *w^1118^* flies. While much research in *D. melanogaster* has used a restricted number of highly inbred or isogenic backgrounds, sleep duration, timing, and architecture vary between *Drosophila* species, as well as inbred, or outbred lines of *D. melanogaster* [49–51]. Artificial selection experiments, phenotyping genetic reference panels, and comparing sleep in across different wild-caught populations reveal genetic background has effects on sleep that are equivalent to those observed in genetically-induced sleep mutants.[52–54]. Analysis of interactions between starvation resistance and the effects of temperature on longevity genetically variable panels suggest that genetic background contributes to interactions between nutrient or temperature stress and longevity [55,56]. While these panels provide a rich resource for examining interactions between sleep and longevity, the characterization of sleep typically focuses on a single time point, limiting the ability to study the relationship between ontogenetic changes in sleep and longevity. Defining the relationship between sleep and longevity across genetically variable populations has potential to inform our understanding of the genetic and functional relationship between sleep and aging.

There is a large body of work in *Drosophila* that aligns with human public health data linking sleep and longevity [25]. Experimental evidence from chronic sleep deprivation studies in both rats and fruit flies demonstrates reduced lifespan [57–59]. However, other studies challenge this connection, showing that long-term sleep deprivation does not necessarily lead to lethality[60]. Genetic studies add further nuance. For example, several short-sleeping *Drosophila* mutants exhibit reduced longevity [25,61,62], whereas mutants such as *fumin*, which lack the dopamine transporter and exhibit severe sleep loss, do not show reduced lifespan [63]. Artificial selection experiments reveal that selecting for long-and short-sleeping flies does not necessarily alter lifespan, suggesting these traits may be genetically and experimentally separable [64]. Taken together, these findings highlight a complex relationship between sleep and longevity, indicating that both the timing and genetic basis of sleep loss are critical factors influencing lifespan.

A central question raised by these studies is the biological significance of the 4–13 day window identified as mediating the effects of sleep on longevity. Prior work has shown that sleep deprivation the first week of adulthood can produce long-lasting consequences, suggesting that neurodevelopment may extend beyond eclosion [28,29]. Sleep is known to influence synaptic connectivity, and age-related changes occur in both central brain neurons and peripheral sensory circuits across the lifespan[65–69]. Although the current studies focused on sleep during adulthood, it remains plausible that sleep earlier in development exerts lasting effects on physiology and behavior. In addition to its neural functions, sleep also regulates peripheral systems, including gut function, adipose tissue homeostasis, and whole-body metabolic rate[70]. Disruption of sleep during this critical window may therefore alter systemic processes in a way that affects long-term health and survival. For example, the transcription factor *pdm3* promotes early life sleep, and it is possible this contributes to long-term changes in sleep and longevity [71]. Identifying this window offers a valuable framework for dissecting the cell-type-specific roles of sleep during early life and how these roles contribute to aging.

Here, we show that pharmacologically increasing sleep by feeding flies with gaboxadol early in development leads to increased longevity. While gaboxadol has been widely used as a sleep-promoting agent, there is evidence that gaboxadol-induced sleep is functionally distinct from natural sleep [72]. Alternative approaches to manipulate sleep include thermogenetic or optogenetic activation of sleep-promoting circuits [44,73,74], exposure to low-frequency mechanosensory stimulation [75], and overexpression of genes that promote sleep [76,77] . However, each of these methods presents limitations, particularly when applied chronically over multiple days. Examining how increasing sleep during early development impacts longevity will be critical for defining the generalizability of the effects reported here.

Longitudinal analysis of behavior has revealed novel interactions between sleep and feeding. These relationships can be further validated using genetic or pharmacological approaches to manipulate specific behaviors. These findings underscore the value of examining how sleep-related genes influence behavior across different life stages, and how these changes contribute to comorbidities. This approach provides a powerful framework for understanding the temporal dynamics of sleep and its broader impact on health.

## Materials and Methods

### Fly Husbandry

All experiments were conducted using *Drosophila melanogaster*. Flies were reared on a standard Drosophila medium prepared according to the Bloomington Stock Center recipe (Bloomington Formulation, Nutri-fly, #66-113, Genesee Scientific, San Diego, California). Flies were maintained in temperature-controlled incubators (Powers Scientific, Warminster, Pennsylvania) at 25 °C under a 12 h light:12 h dark (LD) cycle with humidity maintained between 55–65%. Standard husbandry practices were followed, with all stocks maintained in uncrowded vials and transferred to fresh food every 2–3 days to avoid overcrowding and nutritional depletion. Unless otherwise specified, 3–5-day-old mated females were used for all experiments. Mated females were obtained by maintaining mixed-sex populations following eclosion for at least 48hr. For aging experiments, flies were continuously maintained on standard food and transferred to fresh vials every other day until the desired experimental age was reached. In sucrose supplementation experiments, flies were maintained on standard Bloomington food supplemented with 10% sucrose, unless noted otherwise.

Fly Stocks

The wild-type fly line used in this study was *w^1118^*(Bloomington Drosophila Stock Center, stock #5905) [78]. This line was used for all the experiments. Unless otherwise indicated, experimental manipulations and comparisons were performed relative to this stock. Stocks were expanded and maintained for at least two generations in our laboratory before inclusion in experiments, ensuring adaptation to our husbandry conditions.

### Sleep Behavior Assay

Sleep behavior was measured using the Drosophila Activity Monitoring System (DAM2; TriKinetics Inc., Waltham, MA), which detects locomotor activity based on infrared beam crossings of individual flies. Flies were briefly anesthetized with CO₂ and loaded into 65 mm × 5 mm glass locomotor tubes containing standard Bloomington medium (Nutri-Fly #66-113, Genesee Scientific) or the specified dietary intervention (e.g., sucrose-supplemented food). Flies were acclimated for a minimum of 24 hours prior to the start of behavioral analysis. Food filled at one end of each tube and was sealed with cotton plugs.

Unless otherwise noted, all experiments were performed with 3–5-day-old mated female flies, which were selected to minimize age- and sex-dependent variability in sleep behavior. For lifespan-associated studies, sleep was assayed across different ages, with flies maintained on standard food and transferred to fresh tubes every 5 days to prevent food desiccation and microbial contamination.

For experiments using the Drosophila Activity Monitoring (DAM) system (TriKinetics, Waltham, MA, USA), sleep and waking activity were quantified from infrared beam crossings of individual flies. Raw activity files were retrieved using DAMFileScan (TriKinetics), and custom Python scripts were used to bin activity at 5-min resolution. Sleep was defined as periods of immobility lasting ≥5 min, and sleep traits—including total sleep, mean bout length, bout number, and waking activity—were extracted using the Drosophila Sleep Counting Macro [79–81].

### Longevity Assay

Lifespan was quantified using the Drosophila Activity Monitoring System (DAM2; TriKinetics Inc.), which continuously records infrared beam crossings of individual flies. Flies were collected within 24 h of eclosion and maintained in mixed-sex groups for 2 days to allow mating. Female flies were then separated under brief CO₂ anesthesia and individually loaded into 65 mm × 5 mm glass tubes containing standard Bloomington fly-food (Nutri-Fly, Genesee Scientific). Tubes were maintained in Percival incubators (DR-36VL, Percival Scientific) under controlled environmental conditions (25 °C, 55–65% humidity, 12:12 h light-dark cycle). Flies were transferred to new tubes with fresh food every 5 days, at which point survival was scored. Flies that escaped or were injured during transfers were excluded from the analysis that accounted for less than 5%. The time of death for each fly was operationally defined as the point of last recorded waking activity (final beam break) in the DAM system. Lifespan was calculated as the number of days that survived post-eclosion until this endpoint.

### Temperature and Diet Assays

For these experiments all flies were grown on a standard diet of g/L Nutri-Fly® Bloomington Ready-Mix (yellow cornmeal, agar, corn syrup solids, inactive nutritional yeast, soy flour); 1.0 L DI water; 4.8 mL propionic acid; 10 mL 10% tegosept. For the high sucrose diet, flies were transferred to standard food, supplemented with 10% (w/v) sucrose at 04 days and maintained on this food for the duration of their lives. For high temperature manipulations, flies were grown on standard food at 25°C and transferred to 29°C at 04 days. For the pharmacological induction of sleep, standard food was supplemented with Gaboxadol (THIP): Gaboxadol hydrochloride (T101, Sigma-Aldrich; ≥98% purity by HPLC; synonym: 4,5,6,7-Tetrahydroisoxazolo[5,4-c] pyridin-3-ol hydrochloride) at a final concentration of 0.1 mg/mL on the days specified [82,83].

### Statistical Analysis

All statistical analyses were performed using GraphPad Prism v10.5.0, R (v4.3.2) using RStudio (v2023.06.1+524; Posit Software, Boston, MA). Descriptive statistics (mean, standard deviation, standard error of mean, and range) were generated for grouped datasets. Sleep across aging was analyzed using a second-order polynomial (quadratic) regression model to capture non-linear patterns in lifespan-associated changes, with goodness-of-fit assessed via *R²* values. Group comparisons were evaluated using two-way ANOVA (row and column factors), mixed-effects models (REML) to account for repeated measures, and multiple comparisons testing where appropriate. Survival and longevity analyses were conducted in Prism using the Kaplan–Meier method with log-rank (Mantel–Cox), Gehan–Breslow–Wilcoxon, and Log-rank test for trend as recommended. Simple linear regression was additionally applied to evaluate associations under different experimental conditions (e.g., sucrose vs. temperature stress).

Trait–longevity associations were further plotted using RStudio. Pairwise correlations between sleep traits and longevity were computed using Pearson’s correlation coefficient (*r*) with corresponding 95% confidence intervals, *R²* values, and two-tailed significance tests (α = 0.05). Traits were visualized using bar plots annotated with significance markers, where positive and negative correlations were highlighted in green and red, respectively. Principal component analysis (PCA) was performed on early, mid, and late-life trait datasets after standardization. Longevity was projected as a variable to assess its alignment with trait-based variance structure. Loadings, variance explained, scree plots, and cosine similarity between trait vectors and the longevity projection were calculated to identify the top contributors to each principal component.

Statistical definitions of sleep and activity were based on the Drosophila Activity Monitoring (DAM) system standard, in which ≥5 min of immobility is considered a sleep bout [79–81].

## Acknowledgements

The authors are grateful to Hugo Moebel (TAMU) for technical support. This work was supported by a Texas A&M College of Art & Science Seed Grant to ACK and JB and NIH Grant R01 NS131628 to ACK.

## Supplemental Figures

**Supplemental Figure 1.**
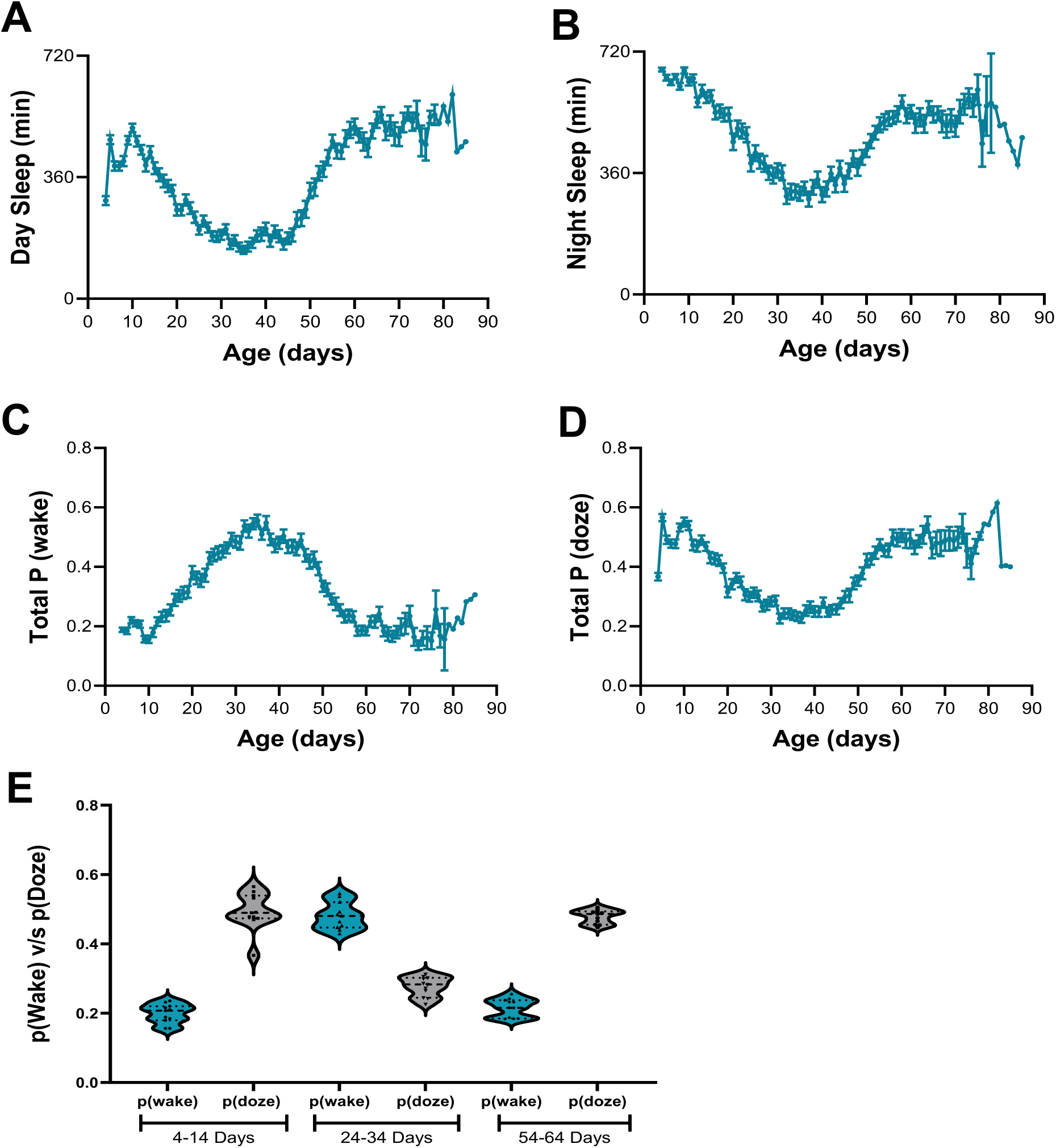
Dynamics of Age-dependent sleep architecture and arousal. S1A) Day Sleep decreased progressively during early life, reached the lowest in mid-life (∼Day 35), and subsequently recovered in late life exhibiting a U-shaped trajectory across the lifespan (*R²* = 0.29). S1B) Night Sleep declined progressively during early life, reached a minimum in mid-life, and recovered partially in late life. A second-order polynomial regression model captured this trend (*R²* = 0.25). S1C) Total probability of wakefulness (P(wake)) increased from early life to a mid-life peak and subsequently declined in older flies (*R²* = 0.29); both linear (95% CI: 0.02251 to 0.02468) and quadratic (95% CI: −0.0003369 to −0.0003074) terms were statistically significant. S1D) Dozing probability decreased during mid-life and recovered old age. A second-order polynomial regression revealed a U-shaped trajectory in dozing probability (*R²* = 0.23). Both the linear (95% CI: −0.01940 to −0.01741) and quadratic (95% CI: 0.0002349 to 0.0002619) terms were statistically significant. S1E) Tukey’s post-hoc test showed strong differences between P(wake) and P(doze) within each age group, including 4-14 days (adjusted p<0.001). Error bars represent SEM at each time point.

**Supplemental Figure 2.**
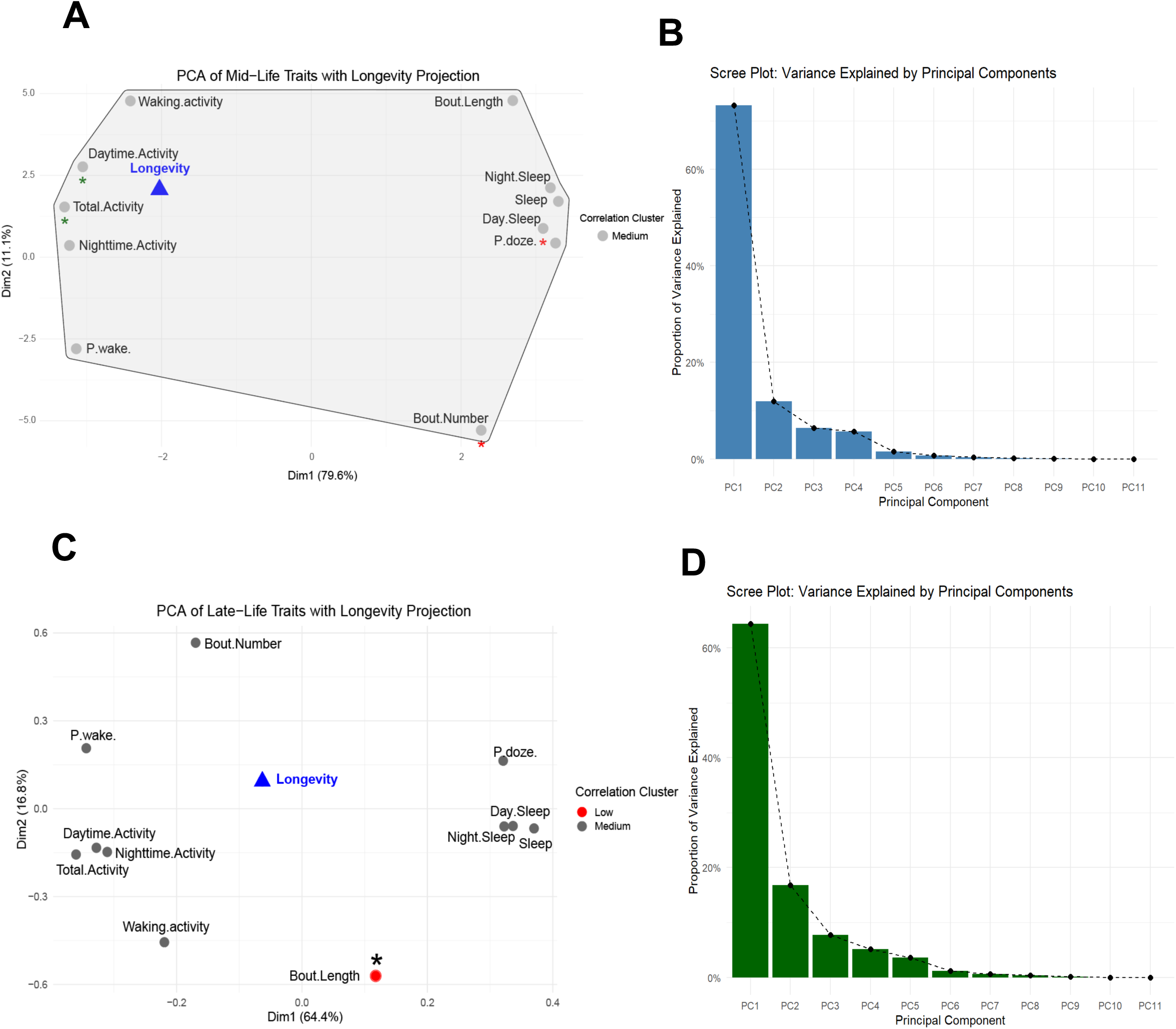
Principal component analysis of sleep and activity traits across life stages in relation to longevity. S2A-B) PC1 separated sleep-related traits (Sleep: 0.33; Doze: 0.33; Night Sleep: 0.32) from activity-related traits (Total Activity: −0.33; Nighttime Activity: −0.32; Wake: −0.31), while PC2 was driven by Bout Number (−0.53), Bout Length (0.48), and Waking Activity (0.48). Traits positively correlated with longevity included Daytime Activity (*r* = 0.25) and Total Activity (*r* = 0.23), whereas Sleep (*r* = −0.17), Doze (*r* = −0.20), and Day Sleep (*r* = −0.22) showed negative associations, indicating that higher mid-life activity and reduced sleep fragmentation predict longer lifespan. S2C-D) PC1 captured variation from reduced Sleep (−0.37), P(doze) (−0.34), and Bout Length (−0.33), while PC2 was shaped by Waking Activity (0.54), Nighttime Activity (0.52), and Total Activity (0.42). However, trait-longevity correlations were weak (e.g., Sleep *r* = −0.05; Activity *r* = 0.13), suggesting that late-life traits poorly predict lifespan. The scree plot (SF2D) shows PC1 and PC2 explain most variance (59%, (PC1 = 42%, PC2 = 17%)), yet this variation is not strongly aligned with longevity outcomes.

**Supplemental Figure 3.**
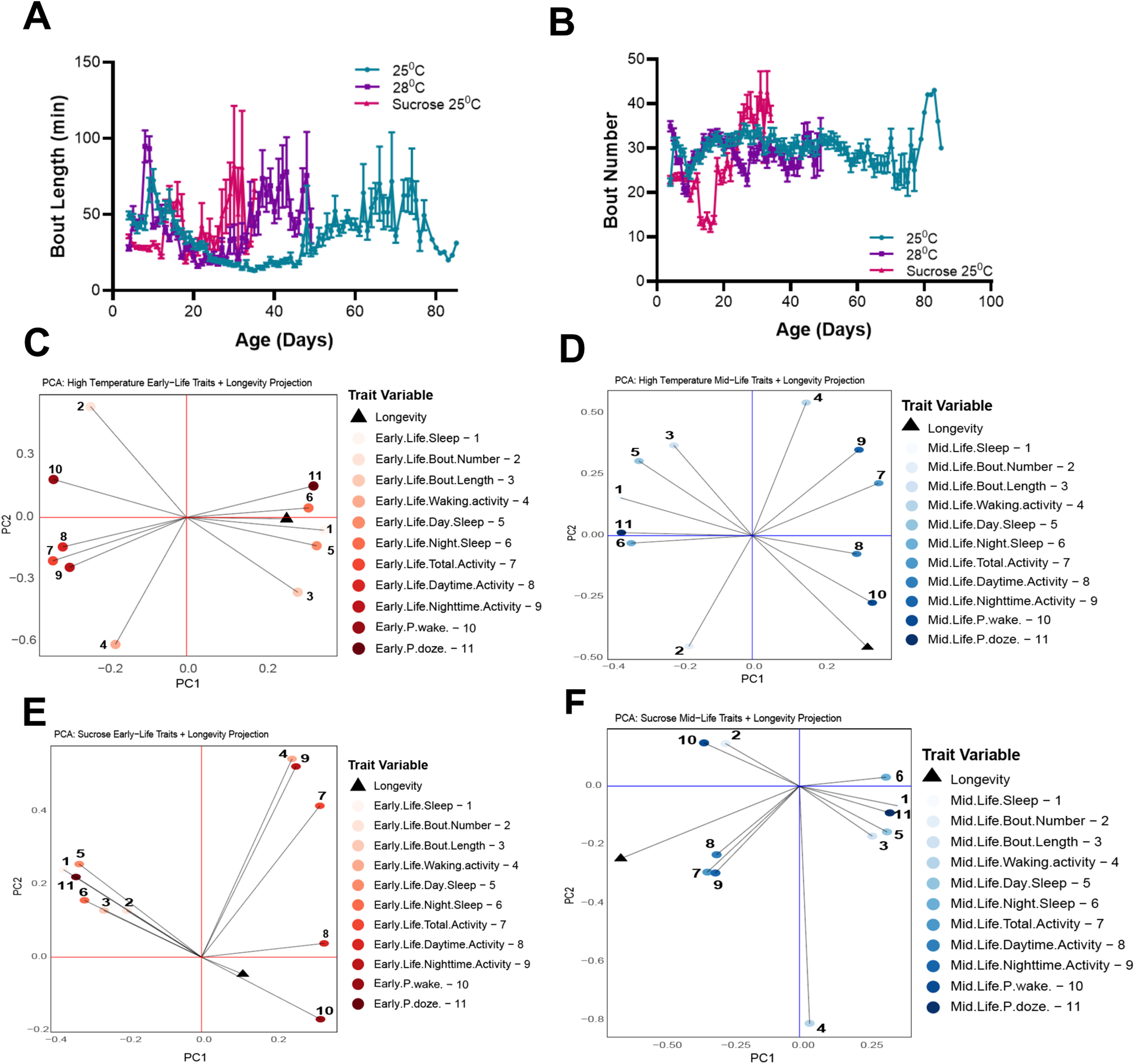
Effect of temperature and nutritional stress on sleep dynamics across the lifespan. S3A) One-way ANOVA revealed no statistically significant difference among groups (*F*(2,155) = 2.665, p>0.05, *R²* = 0.03). Although flies maintained at 25°C lived longer than those at 28°C, the sleep bout length trajectories between these two groups were notably similar throughout the lifespan. S3B) One-way ANOVA revealed a significant main effect of treatment (*F*(2,155) = 4.528, *p*<0.01, *R²* = 0.06). Post-hoc Tukey’s test showed a significant reduction in sleep bout number in the sucrose group compared to control (*p*<0.01), indicating increased sleep consolidation. Although the sucrose group exhibited fewer sleep bouts, their survival was markedly reduced. Notably, flies at 25°C maintained stable bout numbers throughout lifespan, whereas flies at 28°C showed a similar activity pattern despite being short-lived. No significant differences were observed between 25°C and 28°C *(p>0.05)* or between 28°C and sucrose (*p>0.05*). S3C-D) Under thermal stress, early-life PC1 loaded positively on Sleep (0.35), Day Sleep (0.33), and P(doze) (0.32), and negatively on Total Activity (−0.34) and P(wake) (−0.33), while PC2 was shaped by Waking Activity (−0.61), Bout Number (0.54), and Bout Length (−0.36). Sleep showed a positive correlation with lifespan (r = 0.36), while Wake showed a negative correlation (r = −0.26). In mid-life, Bout Length (r = −0.53), Day Sleep (r = −0.42), and Sleep (r = −0.41) showed negative correlations with lifespan. S3E-F) Under dietary stress (10% sucrose), early-life traits showed no correlations with longevity (e.g., Sleep r = −0.11), with PC1 separating rest-related traits (Sleep, P(doze)) from activity (P(wake), Total Activity). In contrast, mid-life traits exhibited negative associations, with Night Sleep (r = −0.64) and P(doze) (r = −0.55), and Activity (r = 0.70), Nighttime Activity (r=0.69) positively correlated with lifespan.

**Supplemental Figure 4.**
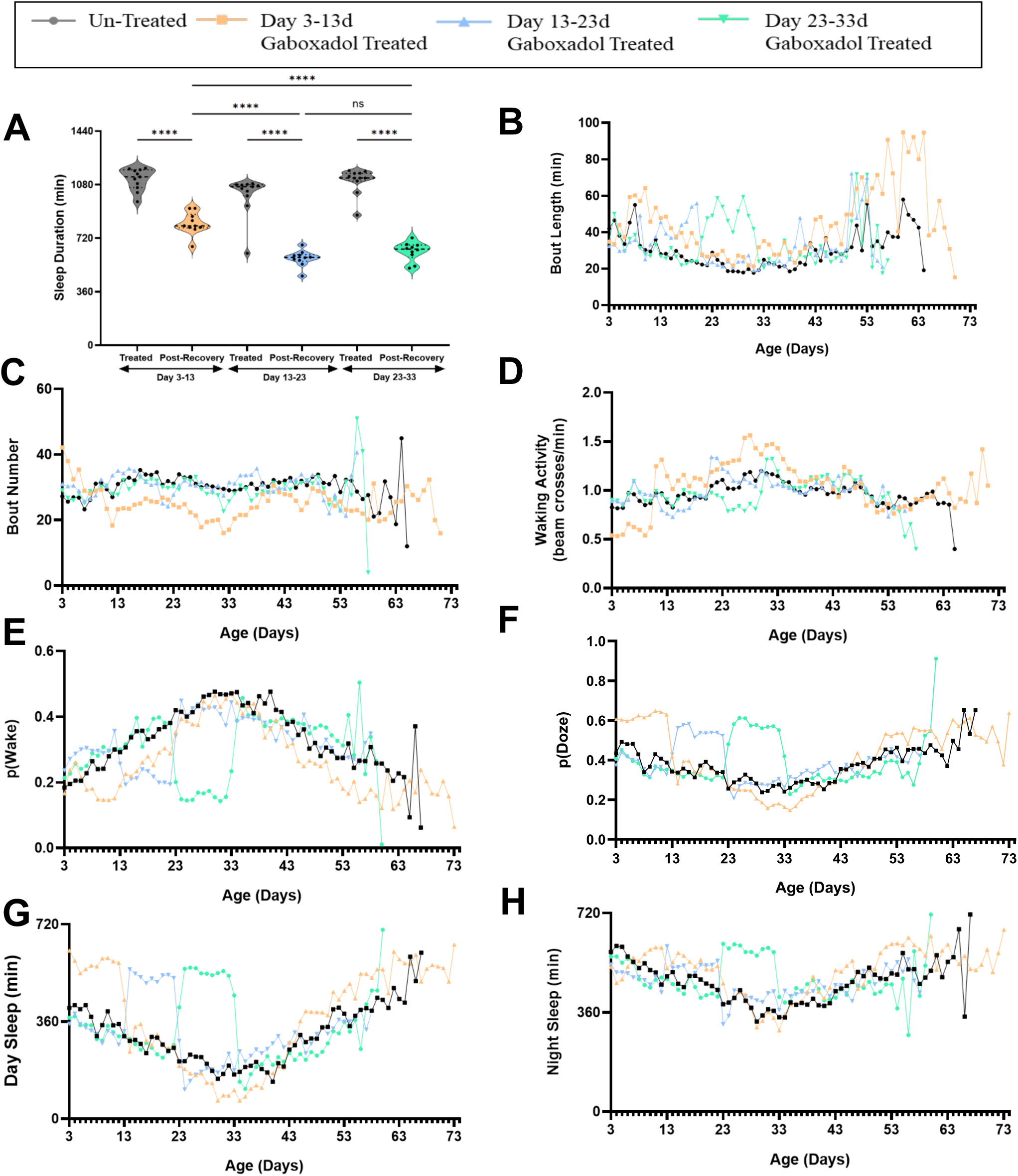
Sleep architecture across lifespan in flies treated with Gaboxadol. S4A) Two-way ANOVA revealed significant effects of treatment window (*F*₁₀,₄₇ = 7.30, *p* < 0.001) and recovery (*F*₅,₄₇ = 172.1, *p* < 0.001). Post hoc comparisons indicated that early-life treatment led to sustained increases in sleep even after drug withdrawal, whereas mid- and late-life treatments did not produce long-term sleep effects. S4B-D) Early-life Gaboxadol treatment enhances sleep consolidation and increases waking activity. Also, early life treatment significantly led to longer average bout lengths (mean diff. = −11.07 min; 95% CI: −13.78 to −8.37; *p* < 0.001) and fewer sleep bouts (mean diff. = +4.71; 95% CI: 3.86–5.57; *p* < 0.001), indicating increased sleep consolidation. Two-way ANOVA confirmed significant effects of age (5.25% and 2.39% of variance) and treatment (1.16% and 2.64%) on bout length and number. A significant increase in waking activity was observed (*p* < 0.001), with age and treatment accounting for 12.8% and 0.6% of the variance. S4E-H) Early-life treatment exhibited strongest association for both P(doze) (*R² = 0.40*) and day sleep (*R² = 0.47*), suggesting improved sleep consolidation. Later treatment windows (Days 13-23 and 23-33) showed lower fit values (*R²* < 0.06).

